# Precise and ultrafast tandem repeat variant detection in massively parallel sequencing reads

**DOI:** 10.1101/2023.02.15.528687

**Authors:** Xuewen Wang, Meng Huang, Bruce Budowle, Jianye Ge

**Affiliations:** Center for Human Identification, University of North Texas, Health Science Center, Fort Worth, TX, USA; Department of Microbiology, Immunology, and Genetics, University of North Texas Health Science Center, Fort Worth, TX, USA

**Keywords:** Tandem Repeat, Variant caller, Long-read sequencing, Whole genome sequencing, Bioinformatic tool

## Abstract

Calling tandem repeat (TR) variants from DNA sequences is of both theoretical and practical significance. A large number of software tools have been developed for detecting TRs. However, little study has been done to detect TR alleles from long-read sequences, and the effectiveness of detecting TR alleles from whole genome sequence (WGS) data still needs to be improved. Herein, a novel algorithm is described to retrieve TR regions from sequence alignment, and a software program, TRcaller, has been developed to call TR alleles from both short- and long-read sequences, both whole genome and targeted sequences generated from multiple sequencing platforms. The results showed that TRcaller could provide substantially higher accuracy in detecting TR alleles with magnitudes faster than the mainstream software tools. TRcaller is able to facilitate scalable, accurate, and ultrafast TR allele calling from large-scale sequence datasets in various applications, such as DNA forensics, medical research, disease diagnosis, evolution, and breeding programs.

**Availability:** TRcaller is available at www.trcaller.com.

## Introduction

The detection of genomic variants is the foundation of most genomic research and applications. These DNA sequence variants mostly include single nucleotide polymorphisms (SNPs), small insertions and deletions (InDels), tandem repeats (TRs), and large structure variations (Willems et al. 2017; Byrska-Bishop et al. 2021). TRs, including both short TRs (STRs), also known as microsatellites, and variable number tandem repeats (VNTRs) or minisatellites, are repeat sequences comprised of a few to many tandem repeat units or motifs. In particular, STRs usually contain repeat sequences that ≤ 6 base pairs (bp) in length, are widely dispersed in genomes, and compose up to ~1-3% of most genomes (Frazer et al. 2009; Chaisson et al. 2015; Wang & Wang 2016). Due to their high variability and discrimination power, TRs have been widely used in forensic identification, studies on species evolution, breeding selection, trait association, clinical diagnostics, medicine design, genealogy, disease diagnosis, and molecular marker development (Frazer *et al.* 2009; Wang & Wang 2016; Saini et al. 2018; Eichler 2019; Chiu et al. 2021). Some TRs are associated with or causative of diseases (Tang et al. 2017; Eichler 2019; Chintalaphani et al. 2021; Depienne & Mandel 2021; Mukamel et al. 2021; Rajan-Babu et al. 2021; Erwin et al. 2022), and detecting such disease-associated TRs and in some situations could provide higher resolving power than SNPs (Saini *et al.* 2018). STRs also are the core markers of most DNA forensic applications and are used in almost all forensic DNA databases, such as the FBI’s Combined DNA Index System (CODIS) database (fbi.gov 2022).

Traditionally, DNA variants have been detected by Sanger sequencing or by measuring the lengths of DNA fragments. The development of massively parallel sequencing (MPS) has augmented detection of DNA variants. The second generation sequencing technologies (e.g., NovaSeq 6000 from Illumina) are able to sequence millions of short DNA fragments simultaneously and produce short reads up to 250 bp, and provides high accuracy of detecting SNPs and other short variants (Stoler & Nekrutenko 2021). STRs may also be detected with the second generation sequencing technologies (Zeng et al. 2015; Churchill et al. 2016). However, sequencing through a long TR region is still challenging with Illumina platforms (Gettings et al. 2019). The third generation single molecule sequencing technologies, such as Oxford Nanopore Technologies (ONT) MinION and Pacific Biosciences (PacBio) Revio, are able to sequence long DNA fragments (1,000s ~ 100,000s bp) and have revolutionized studies in genome assembly, association studies, structure variant detection, etc. (Chaisson *et al.* 2015; Logsdon et al. 2020). Particularly, with the recently developed HiFi method, the accuracies of SNP detection with PacBio single-molecule real-time (SMRT) sequencing have been substantially improved up to 99.8% (Wenger et al. 2019). Additionally, the alleles of long TRs are detectable (Gettings *et al.* 2019).

While the TR regions may be sequenced with MPS technologies, precisely calling the TR alleles still can be challenging. TRs vary among individuals of a population in repeat motif lengths, number of repeats, partial bases of a motif and sequence like SNP (Gymrek et al. 2017). Gaps from contraction or insertion from expansion are common throughout TR regions, which makes alignment within the TR regions problematic. A number of bioinformatics tools have been developed for detecting STR loci and calling the STR alleles in either whole genome sequence (WGS) and targeted sequencing data, including lobSTR (Gymrek et al. 2012), HipSTR (Willems *et al.* 2017), GMATA (Wang & Wang 2016), Tandem Repeat Finder (Benson 1999), STRait Razor (Woerner et al. 2017; King et al. 2021), Universal Analysis Software (UAS, www.illumina.com), Straglr (Chiu *et al.* 2021), RepeatSeq (Highnam et al. 2012), etc. Some tools may even infer STR alleles that are longer than individual reads, such as ExpansionHunter (Dolzhenko et al. 2017; Dolzhenko et al. 2019) and Tredparse (Tang *et al.* 2017). Among those programs, HipSTR was designed for calling STRs from short reads via realigning and optionally eliminating PCR stutters (Willems *et al.* 2017), which was shown to outperform lobSTR in terms of accuracy of calling STR alleles. Straglr was developed to call STR alleles by clustering and statistical modeling from long reads of at least 200 bp (Chiu *et al.* 2021), but not designed for short reads. However, Straglr does not report specific, accurate STR allele sequences, but instead provides a range of STR distributions, which may not be acceptable for applications that require precision allele calling (e.g., forensics). ExpansionHunter uses read reassembly and sequence graph to discover TRs, even longer than read length (Dolzhenko *et al.* 2017; Dolzhenko *et al.* 2019). RepeatSeq uses the GATK tool (Van der Auwera & O’Connor 2020) and statistical models to call STR alleles (Highnam *et al.* 2012). STRait Razor and UAS were specifically designed for forensic applications, which usually accept high coverage (e.g., 10s~1000s) short reads from targeted amplified regions, such as the STR regions included in a MPS based STR kit (e.g., ForenSeq DNA Prep Kit, Verogen) (Alonso et al. 2018). All software programs designed for forensic applications require anchor sequences typically on both sides of the STR regions to identify predetermined markers. With this approach, a relatively high allele dropout rate can occur with typical short-read WGS data with 30x or lower coverage, because the low coverage short reads may not contain both sides of the anchor sequences or may not contain long STR alleles. In addition, these software programs may generate false positive results for long-read sequences, because the same anchor sequences may exist in multiple positions across the genome or even in the same long read. To the best of our knowledge, no software tool has been developed to precisely call TR allele sequences from long-read sequences.

Herein, we describe the development of a software program, TRcaller, that implements a novel algorithm for calling TR allele sequences from both short- and long-read sequences, generated from either whole genome and targeted sequences, and achieves greater accuracy and sensitivity than existing tools. The accuracies of the algorithm were evaluated and compared with those of the major existing software tools using the short and long-read sequences generated from multiple platforms: Illumina HiSeq, NovaSeq, MiSeq, PacBio, ONT, and 10X genomics, as well as with simulated reads. TRcaller can run seamlessly across multiple computer operating systems, and the software program was further optimized to call the TR alleles from a large scale WGS data in seconds with a regular computer.

## Results

### Overview of the workflow and algorithms of TRcaller

Novel algorithms and workflow were developed to analyze sequence data and detect TR alleles efficiently and accurately from both short and long read sequences. The algorithms and workflow were implemented into a software called TRcaller (Figure 1). The workflow takes an aligned sequence file in indexed BAM format (together with a BAI index file) and a target TR loci file in BED format as input, and outputs the TR allele length/size, allele sequences, and supported read counts in the sequence data (Figure 1). The index of alignment was used to quickly locate the targeted TR loci, which enables ultrafast TR analysis (e.g., ~2 seconds for detecting alleles of 20 STR loci in a 300x mean coverage WGS human sample). There are three major modules of the TRcaller workflow, including input, TR allele detection, and output. The input module takes an alignment file in BAM format, together with a BAI index file, and a BED file describing the details of the targeted TR loci. First, in the TR allele detection module, only aligned reads that cover the targeted TR regions are retrieved with the index and locus position information in the BED file. Second, the boundaries of TRs are determined, the TR sequences are extracted with the determined boundaries, and the alleles are called after optional motif validation and assigned into alleles based on sequences. Finally, the output module generates reports.

**Figure 1.**
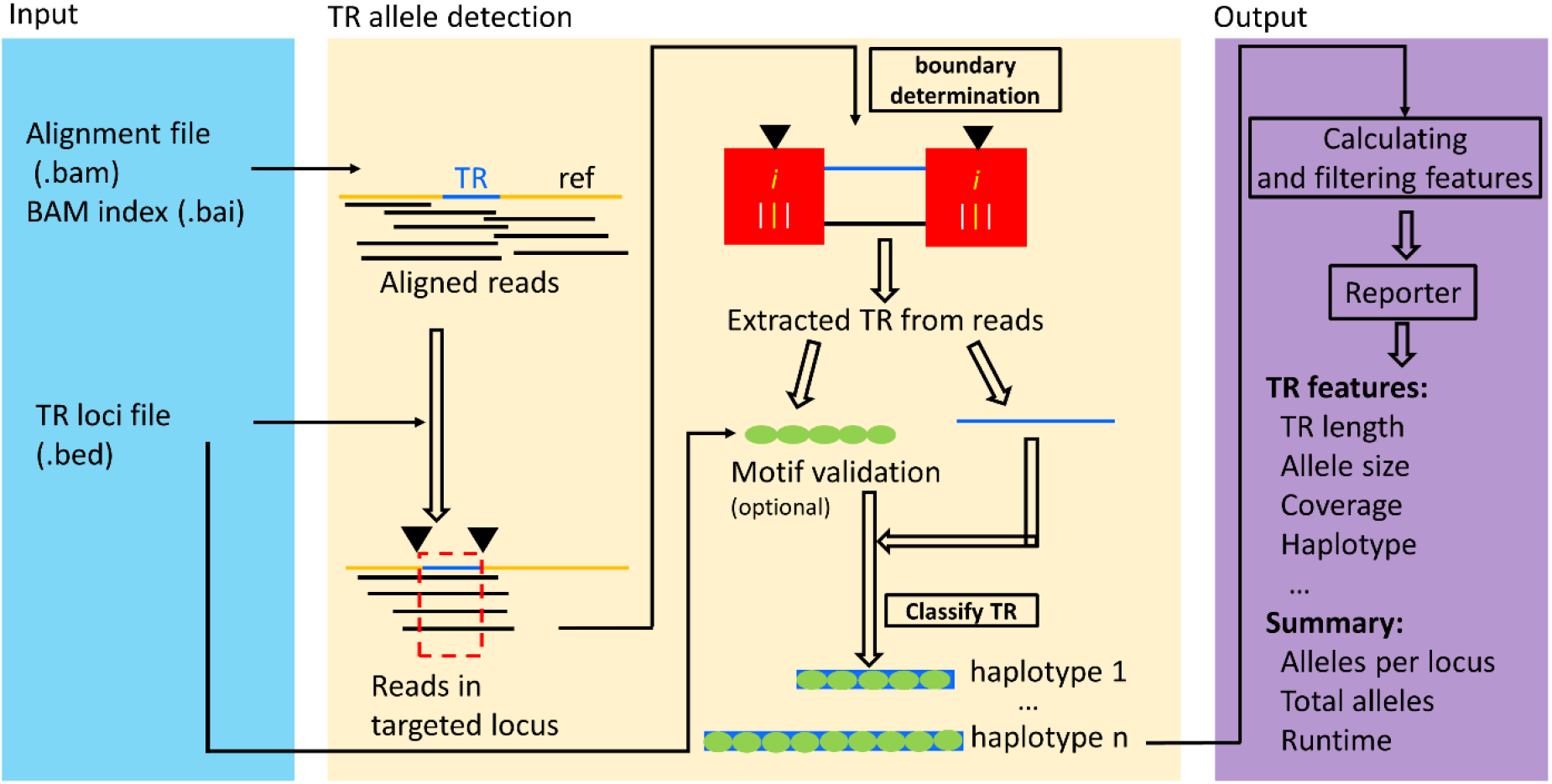
The schematic workflow of TRcaller.

### Sensitivity evaluation of TRcaller via simulated MPS reads

To assess the sensitivity of TRcaller, the calling rate of TR alleles were evaluated from three simulated datasets with different read depth and read length from simulated human genomes with known STR alleles, including Illumina paired-end 150 bp (PE150), Illumina paired-end 250 bp (PE250), and PacBio long CCS (PLCCS) reads for the 20 CODIS core STR loci (Supplementary Table S1) (Hares 2015). At least 100 simulations were conducted at each read depth coverage. The accuracy was calculated as the number of correctly called alleles divided by the expected number of alleles. The allele dropout rate was calculated as the number of correctly called loci divided by the expected number of loci. Incorrectly called alleles were designated as those that did not match the ground truth, which were not observed in the simulated dataset. For all simulations, the detected STR alleles were identical to the expected ground truth (i.e., 100% accuracy) (Figure 2, Supplementary Table S2 and Table S3). However, the average coverage and the average read length can have a substantial impact on the calling rate, particularly when the coverage is low (Figure 2). To reach a 99.9% recalling rate, at least 25x, 10x, and 5x average depths were needed for PE150, PE250, and PLCCS reads, respectively (Figure 2 A and B). These results suggest that longer reads could yield a better TR recalling rate with lower read depth than short reads. In addition, shorter average read length results in a higher allele dropout rate or missing call rate (Figure 2 C and D). In particular, one of the largest amplicon CODIS loci, D21S11, with 127 bp in the reference genome, had a higher allele dropout rate than the other loci at 5x coverage in either PE150 or PE250. In summary, these simulation results suggest that TRcaller can reach a high sensitivity for sequence data with relatively high coverage and/or high average read length.

**Figure 2.**
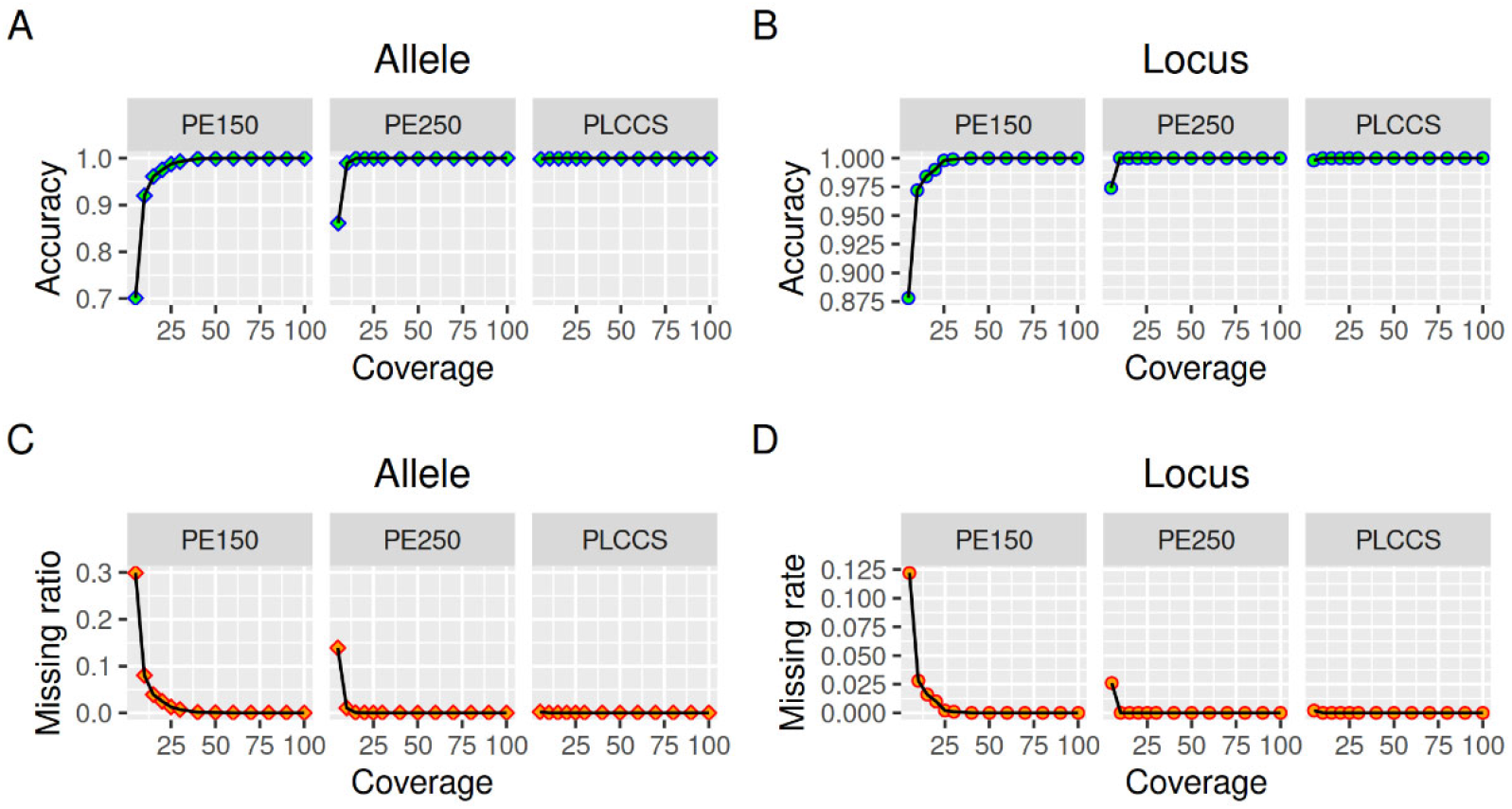
Sensitivity of TRcaller from simulated reads with various coverages and lengths.

### Accuracy evaluation between TRcaller and HipSTR with known profiles

To evaluate the accuracy of TRcaller, 20 CODIS core STR loci from 289 WGS data samples with 30x coverage by the NovaSeq 6000 from the 1000 genomes project (Byrska-Bishop *et al.* 2021) were called using both TRcaller and HipSTR, and the called STR alleles were compared with previously published alleles (Aalbers et al. 2020), in which the sequence data were generated by the Verogen ForenSeq kit, and the alleles were called by Illumina UAS software. The allele identification criteria were set to a minimum coverage of 2x and a maximum two reported alleles for both TRcaller and HipSTR. In total, there were 11,003 alleles reported in the targeted amplification by Aalbers et al. 2020. TRcaller and HipSTR detected 10,310 and 9,946 alleles, respectively (Table 1). Among the detected alleles, 99.38% and 93.42% of alleles from TRcaller and HipSTR, respectively, were consistent with the alleles detected by targeted amplification. The inconsistent alleles had either the same length but different haplotypes (0.20%), or different lengths (0.38%), which most likely are due to amplification and sequencing errors. The alleles that were not detected by TRcaller were mainly due to short-read sequencing that was not able to sequence through large STR alleles, although the average read coverage of the WGS data was about 30x. More details may be found in the supplementary materials (Supplementary Table S4 and Table S5). Overall, these results support that TRcaller can achieve a higher recovery rate from WGS data with >99% accuracy, barring amplification and sequencing errors, than one of the current mainstream software.

**Table 1.** Comparison between TRcaller and HipSTR for calling 20 CODIS STR loci.

| Category | TRcaller |  |  | HipSTR |  |  |
| --- | --- | --- | --- | --- | --- | --- |
| | TRcaller alleles | % true alleles<sup>#</sup> | % called alleles<sup>\$</sup> | HipSTR alleles | % true alleles | % called alleles |
| Alleles consistent with target amplifications | 10,246 | 93.12% | 99.38% | 9,292 | 84.45% | 93.42% |
| Inconsistent alleles |  |  |  |  |  |  |
| Same length but different haplotypes | 22 | 0.20% | 0.21% | 10 | 0.09% | 0.10% |
| Different lengths | 42 | 0.38% | 0.41% | 654 | 5.94% | 6.58% |
| Called alleles | 10,310 | 93.70% | 100.00% | 9,946 | 90.39% | 100.00% |
| Missing alleles | 693 | 6.30% |  | 1,057 | 9.61% |  |
| Total alleles with target amplifications* | 11,003* | 100% |  | 11,003* | 100% |  |

\* the number of true alleles detected with a targeted commercial panel in (Aalbers et al. 2020) with an average coverage of 1,825x.
<sup>#</sup> (Consistent alleles)/(Total alleles)\*100%; <sup>\$</sup> (Consistent alleles)/(Called alleles)\*100

### Cross-platform comparisons of TR calling

To evaluate the performance of TRcaller in calling alleles in sequences generated from various platforms, the data generated for benchmark human reference HG002 in the Genome In A Bottle project (GIAB) (Zook et al. 2016) were used, including both short reads (Illumina NovaSeq WGS 2×250 bp, 10X Genomics Chromium with HiSeq2500, and Illumina NovaSeq WGS 2×150 bp) and long reads from PacBio CCS 15kb_20kb chemistry2 and Oxford Nanopore MinION R10.4 (Supplementary Table S6). In the analysis, the minimum read coverage threshold for calling TR alleles was set at 2x for all platforms, and the minimum allele proportion was set at 0.1.

TRcaller was able to detect all expected alleles at all loci from both PacBio and Illumina 250-bp datasets (Figure 3), which were 100% concordant with the alleles detected with the ForenSeq kit (Gettings *et al.* 2019). All expected alleles also were detected from 300x Illumina 150 bp paired-end reads, except one allele (i.e., 31.2) at the locus D21S11, as this allele is relatively long. Thus, long reads generated more STR results than short reads, even though the short reads data had a higher read coverage. In addition, both alleles at the D21S11 locus were not detected in 10X genomics data, which indicated that the optimized sequencing technology with the same read length might not improve TR allele calling. However, the TR calling accuracy from the Oxford Nanopore MinION data was much lower than data generated on other platforms. In general, TRcaller is capable of accurately calling TR alleles for both short-read and long-read sequences, and longer reads will provide better resources to detect more accurate TR alleles.

**Figure 3.**
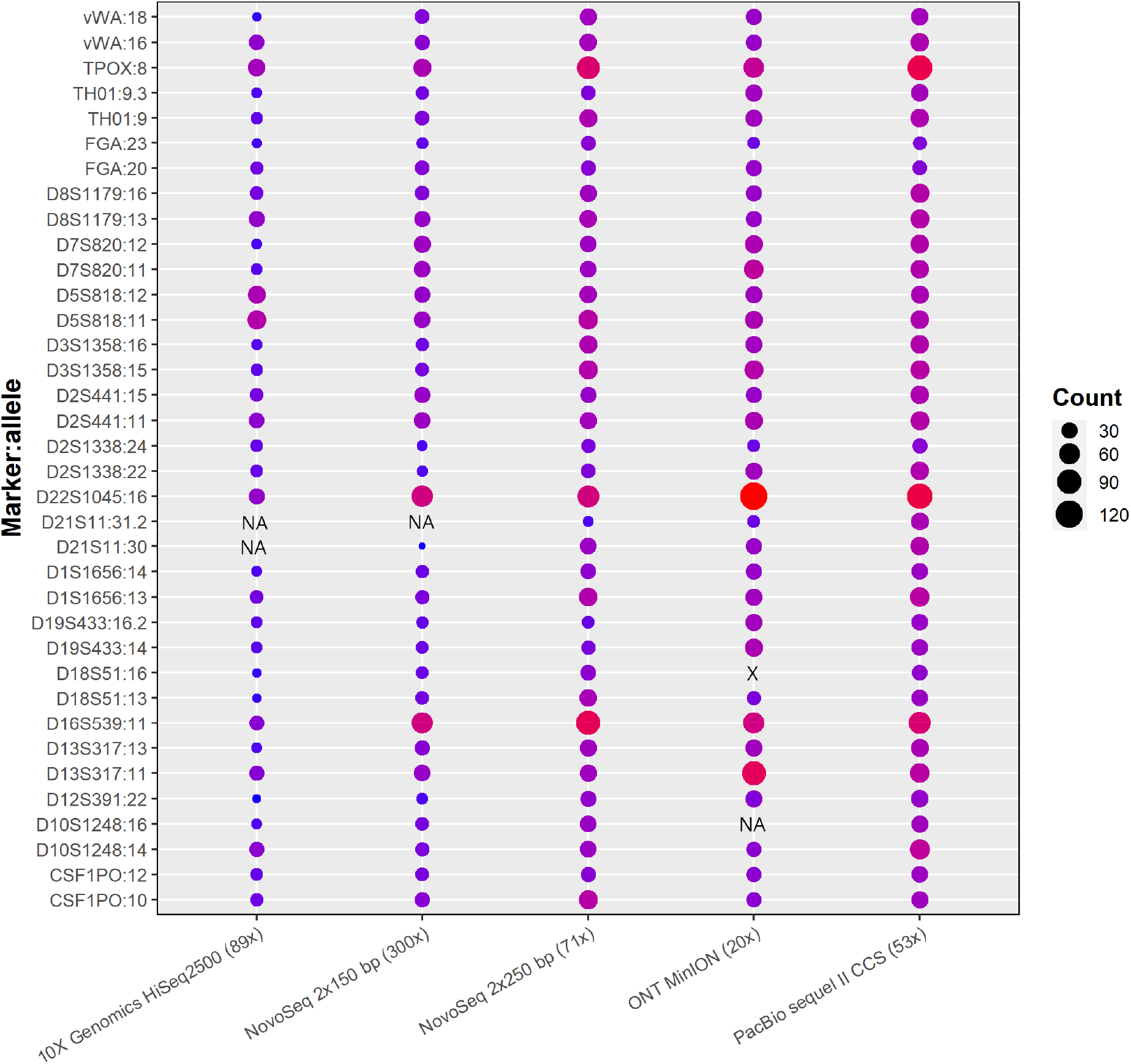
STR alleles called from benchmark human reference HG002 with WGS data generated from various platforms. The number inside parentheses is the average read coverage. The shaded circles represent identical alleles detected in the data set. The plotted read count was after normalization of the original TR allele containing read counts to 100x input. The marker:allele represents the known allele size of each marker (Gettings et al. 2019). NA and X stand for not detected and wrong allele size, respectively.

### Performance of detecting variants in long tandem repeats associated with disease

Sequence data from the Pacbio website was downloaded and used to evaluate the performance of TRcaller for detecting alleles in disease-associated TRs with high numbers of repeats. This dataset contains seven individual samples, generated with PacBio targeted sequencing (Supplementary Table S6 and Table S7), and three autosomal loci *HTT*, *C9orf72*, *ATXn10*, and one X chromosomal locus *FMR1* were included. ATXN10, C9orf72, FMR1, and HTT represent the long tandem repeat loci of gene ATXN10 associated with Parkinson’s disease, gene C9orf71 associated with amyotrophic lateral sclerosis, gene FMR1 associated with fragile X-associated with primary ovarian insufficiency, and gene HTT associated with Huntington disease of human, respectively. In this analysis, the minimum coverage was 2, the minimum allele proportion was 0, and the maximum number of donors was 1. All these four TRs were successfully detected and were 100% concordant with previously analyzed results (Supplementary Table S8). Figure 4 shows the alleles from sample bc1015 with a minimum read depth coverage of 10x, and all alleles were correctly called.

**Figure 4.**
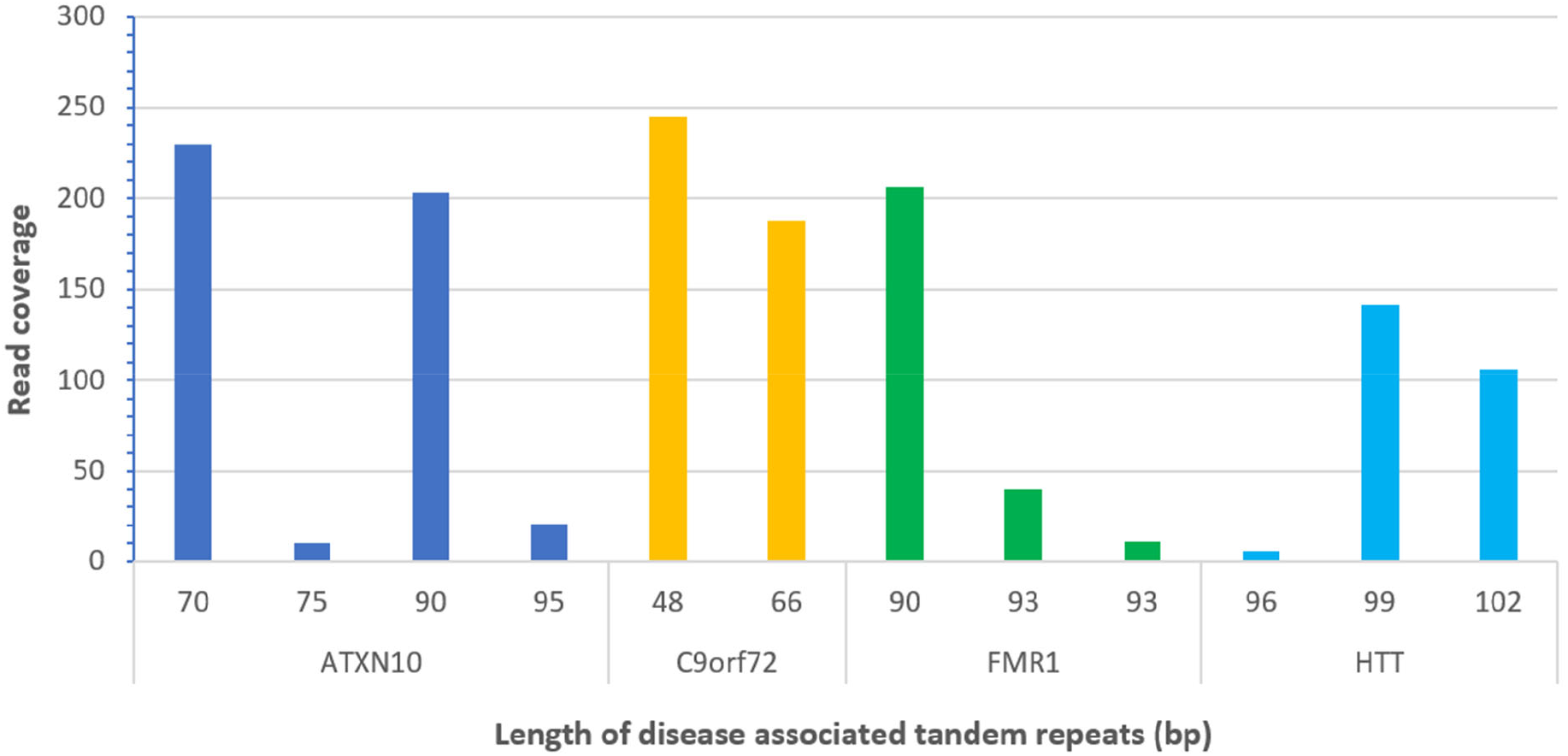
Detected alleles of disease-associated tandem repeats from PacBio long read sequences (sample bc1015).

### Ultrafast STR allele calling and quantification with TRcaller

The processing times of calling TR alleles were compared between TRcaller (v1.0), HipSTR (v0.6.2), and STRait Razor (v3.01). Targeted sequence reads, Illumina WGS reads, and PacBio WGS reads of human samples from the 1000 Genomes Project or the Genome In A Bottle (GIAB) project (Zook *et al.* 2016; Foox et al. 2021) were used to call the alleles of 20 CODIS core STR loci (Hares 2015) (Table S1). Table 2 shows the processing times of calling STR alleles with these three software programs. For the targeted sequence dataset, which is much smaller than the WGS datasets, both TRcaller and STRait Razor required similar processing times that were faster than HipSTR. For WGS datasets, TRcaller only took less than 2 seconds to complete the analysis on either Linux or Window platforms. The speed of TRcaller is multiple magnitudes faster than both STRait Razor and HipSTR, which demonstrates that Trcaller is ultrafast and scalable to large-scale deep sequencing data. HipSTR could not run in Windows, nor process long-read sequence data. STRait Razor worked in Windows, but a large number of incorrect STR alleles were detected in the WGS data, likely because it was not designed for WGS data.

**Table 2.**
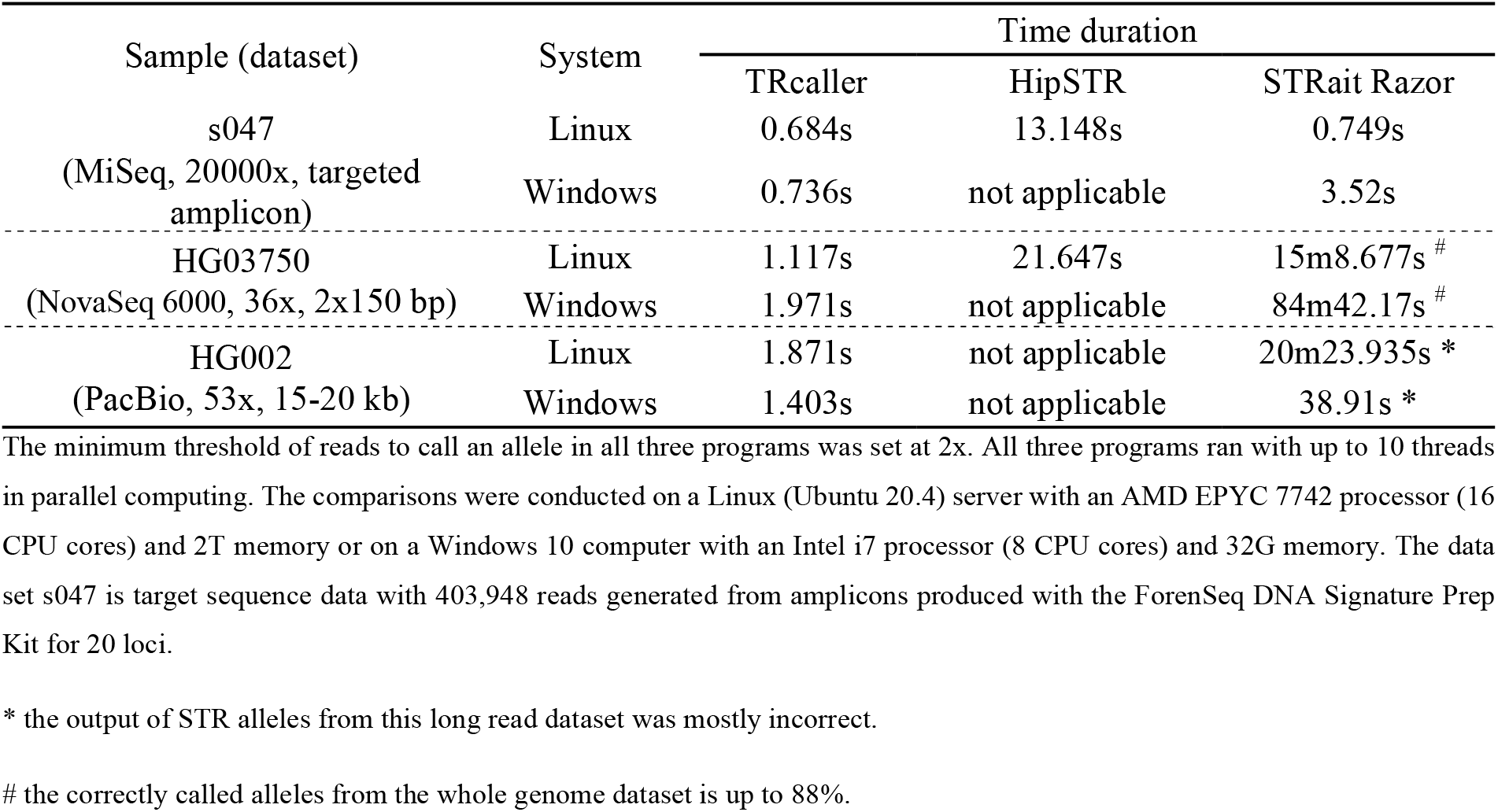
Processing time comparison of three tools.

## Materials and Methods

### The workflow of TRcaller

TRcaller can efficiently call targeted TR alleles from any DNA sequence reads, as long as the alleles with a few base pairs in the flanking regions are fully sequenced in individual reads. Figure 1 shows the schematic workflow of TRcaller. First, TRcaller takes three files as the input: a BAM file, an index BAI file of the BAM, and a BED file. The BAM file contains the alignment data, which may be generated by any mapping tool (e.g., BWA and minimap2) against a reference genome. The BED file contains relevant information about the targeted TR loci, including the name of each TR, the chromosome ID where each TR is located, the start and end positions of each TR, the repeat motif sequences, the length of the repeat motif, and the number of internal offset in base pairs within a TR locus that should be excluded when converting the sequence-based TR alleles to length-based TR alleles, an optional minimum proportion of each TR locus specific stutter ratio threshold, and ploidy.

Second, the boundaries of the TRs are determined by comparing the read mapping locations and the TR regions defined in the BED file. Then, TRcaller extracts the DNA reads that cover the targeted TR regions based on the alignment, which can substantially speed up the analysis time for WGS data since the majority of the reads would not be mapped to the targeted TR regions defined in the BED file. For example, the sequences of the 20 core CODIS STR loci (Hares 2015) are only about 0.0001% of the human genome.

Third, with the boundaries determined, the targeted TR sequences are extracted after trimming the bases outside of the boundaries. These extracted sequences are further compared with the repeat motif sequences in the BED file for allele validation. Next, the same targeted sequences at each TR are phased into haplotypes based on the context of bases, and the coverage of each haplotype is counted by its occurrences. Further, only haplotypes that meet the predefined thresholds may be called as TR alleles, which include the minimum read coverage threshold (e.g., ≥2x for 30x WGS data, or ≥10x for targeted amplification data) and a minimum proportion threshold (e.g., ≥10% in all reads covering a TR locus). TRcaller also requires the user to decide the maximum number of donors or contributors to a sample, which will be used in deciding the maximum number of alleles at a locus. If the DNA sequences are generated with a PCR process, a locus-specific stutter threshold (e.g., 25%) may be applied to filter out the stutter products at each TR locus, but this threshold will only apply to single source samples. For mixture samples, the stutter ratio threshold would not apply. In addition, the maximum possible number of alleles for a locus will be the product of the maximum number of donors and the ploidy of the locus (e.g., a two-person mixture at an autosomal locus can have up to 4 alleles).

Further, for applications that define TR alleles by the number of repeats inferred by the length of amplicon (e.g., the forensic STRs), the sequence-based TR alleles are converted to length-based TR alleles for backward compatibility with allelic data in all national DNA databases worldwide. The approach is based on the recommendations (Alonso *et al.* 2018; Phillips et al. 2018) and formulated with the equations below and adopted from USAT (Wang et al. 2022). Briefly, the length-based TR alleles usually contain both an integer part and a fractional part, separated by a dot (e.g., 5.1). The fraction part may be omitted if it is 0.

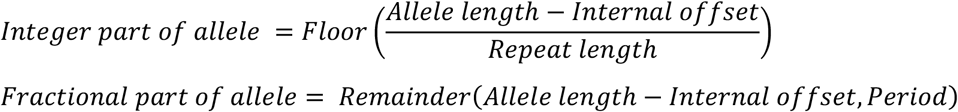

in which the Floor(x) is the function to output the greatest integer less than or equal to x; Remainder(x, y) is the remainder of x divided by y; the allele length is the total number of bases of the allele; the internal offset is the number of bases that need to be excluded in counting the length of alleles, and the repeat length is the length of the repeat motif. For example, for a TR with a motif of ATCG (repeat length = 4) and an internal offset of 2, the integer part of the allele size of a sequence allele “ATCGATCGggATCGA” (“gg” as internal offset sequences) would be Floor((15-2)/4) = 3, and the factional part is the remainder of (15-2)/4, which is 1, and thus the length-based allele size would be 3.1.

Finally, a report of the called TR alleles is generated, which includes the sequence-based TR alleles, length-based TR alleles, the read coverage of each allele, and other relevant summary statistics for each marker. All the thresholds used in the analysis will be included in the final report.

### Software implementation and testing

TRcaller was fully implemented with Java (version 16.0.1) and can be run on any operating system with a JAVA running environment installed. The HTS library java version HTSJDK (version 2.24) was used for the reading and parsing the BAM file (Bonfield et al. 2021). To facilitate the usage of this software, two preprocessing tools were also implemented, one for sorting and indexing a BAM file and one for reducing the BAM file size by only extracting the reads covering the targeted regions. TRcaller has been tested and works well in Windows 10 (version 21H1), MacOS (version 11.6), and Ubuntu Linux (version 20.4), and is currently hosted at www.trcaller.com.

### Read simulation and sensitivity analyses

To test the sensitivity of TRcaller, the calling rates for each allele and each marker locus were evaluated with various sequence read lengths and depth coverages. WGS DNA sequences were randomly simulated from two genomes, the human reference genome GRCh38 and an alternative simulated genome with known mutated TR variants while other sequences are identical to GRCh38, for the 20 CODIS core STR loci (Supplementary Table S1), with a series of average read depths of 5x, 10x, 15x, 20x, 25x, 30x, 40x, 50x, 60x, 70x, 80x, 90x, and ~100x. The coordinates of simulated reads are known, and thus the true TR alleles of these datasets are known. The paired-end 150-bp and 250-bp were simulated with the Wgsim (Danecek et al. 2021), and PacBio long CCS reads were simulated with a 95% accuracy with the Simlord (Stöcker et al. 2016). For each coverage, 100 rounds of simulations were performed. TR alleles were called with TRcaller and then compared with the ground truth alleles. Distributions of sensitivity were plotted with the ggplot2 package for R language (version 4.1.2) (Wickham 2016).

## Discussion

Calling TR variants from DNA sequences is more challenging than calling SNPs due to the large size of the alleles and the inherently complex and variable nature of highly repeatable regions within and in near flanking regions. The current mainstream software programs for calling STR alleles, such as HipSTR (Willems *et al.* 2017) and Strait Razor (Woerner *et al.* 2017; King *et al.*2021), were designed for short reads. The software package TRcaller reported here was developed for calling TR variants from both short and long reads, generated either by whole genome or targeted sequencing. The outperformance in accuracy, speed, and scalability of TRcaller over HipSTR and Strait Razor is the result of the novel data filtering strategy, which uses the indexed genomic mapping information from a binary alignment file to directly locate target locus-based reads by precisely determine the TR boundaries, thus substantially reducing the number of reads needed to be analyzed. It enables an efficient solution for TR haplotype calling from any sequence data. It is worth pointing out that TRcaller uses an alignment strategy to define the boundaries of TRs; in contrast, most existing similar application tools use either the reference genome sequence or anchor sequences in the flanking regions. Mutations in the flanking region may have more effects on the calling accuracy for the strategy with anchor sequences.

The success rate of calling TR alleles can be affected by the read length (i.e., sequencing technology) and the read depth coverage. Large TR loci (e.g., D21S11) tend to have higher allele dropout rates, compared with the small TR loci. Higher read depth coverages can reduce the dropout rates, but cannot overcome the technology limitation regarding read length (e.g., < 250 bp). With long-read sequencing technology, a read depth coverage as low as 5x can recover almost 100% of TR alleles. As long as the quality and quality of template DNA are sufficient, PacBio HiFi reads performed the best of the methods tested herein for TR calling. Based on this study, PE250 and long-read sequencing are recommended for a high-quality TR calling.

The computing time reported in Table 2 did not include the time for generating alignment in BAM format for TRcaller and HipSTR (Willems *et al.* 2017). However, the indexed BAM files are widely accepted as input in other variant callers, such as GATK (Van der Auwera & O’Connor 2020), HipSTR (Willems *et al.* 2017), and DeepVariant (Yun et al. 2021). For the alignment data tested, TRcaller accepts alignment files generated from a wide range of read aligners used in the GIAB project, including BWA (Li & Durbin 2010), novoAlign (http://www.novocraft.com/), (Raczy et al. 2013), minimap2 (Li 2018), and pbmm2 in the package SMRTlink (version 10, https://www.pacb.com/), etc. Therefore, TRcaller should be able to accept generic alignment in BAM format generated from most, if not all, aligners.

Stutters generated during the PCR process are common artifacts that can be detected by TR variant callers (Hoogenboom et al. 2017; King *et al.* 2021). TRcaller uses a minimum coverage and a minimum stutter ratio threshold for each locus to filter out the noisy stutters for single source samples. In the tests with CODIS loci from both whole genome and targeted sequence datasets, a minimum read depth coverage of 2x and a minimum stutter ratio threshold of 0.25 were sufficient to filter out stutters and maintain accuracy.

One limitation of TRcaller is that it assumes the completeness of the TR allele in a sequence read. If the TR allele is not fully present in a read, the read will be skipped by TRcaller, although some tools use an assembly strategy and statistical probability to infer the TR allele, such as Tredparse (Tang et al. 2017) and ExpansionHunter (Dolzhenko *et al.* 2017). From this aspect, TRcaller detects the evidence-based TR alleles with high precision. In addition, TRcaller requires aligned sequences to detect alleles, which is different from some tools (e.g., STRait Razor) that can directly detect TR alleles from FASTQ files.

In summary, TRcaller is a novel software to facilitate scalable, accurate, and ultrafast TR allele calling from large scale sequence datasets in various applications, such as forensics, medical research, disease diagnosis, clinical testing, evolution, and breeding programs. Additionally, the output from TRcaller provides all the information which meets the latest requirements of forensic STR submissions into the databases CODIS or STRidER (Bodner et al. 2016).

## Acknowledgment

The authors would like to thank Katherine Gettings and Sanne Aalbers for sharing their data.

## Supplementary file

Table S1. Forensic CODIS core STR loci and configuration information for the human genome

Table S2. Statistical summary of TRcaller called alleles from simulated read datasets

Table S3. Statistical summary of TRcaller called marker loci from simulated read datasets

Table S4. The comparison of STR alleles between published data from Aalbers et al. 2020 and HipSTR

Table S5. The comparison of STR alleles between published data from Aalbers et al. 2020 and TRcaller

Table S6. Links to publicly available sequencing datasets used in this study

Table S7. Configuration information of 4 disease-associated TR loci

Table S8. The previously identified alleles from disease-associated tandem repeat loci

## Contributions

Jianye Ge developed the algorithm and completed the first version of the software. Xuewen Wang further refined the algorithm and optimized the parameters. Meng Huang tested the software. Xuewen Wang prepared the first draft. Jianye Ge and Bruce Budowle reviewed and revised the manuscript. All authors read and approved the final version of the manuscript.

