## Supplementary data for "Precise and ultrafast tandem repeat variant detection in massively parallel sequencing reads"

### Table S1. Forensic CODIS core STR loci and configuration information for the human genome

| Chrom | ChromStart | ChromEnd | Name | Repeat length | Motif | Ref_hap_length | Ref_allele | Stutter_ratio _threshold | Ploidy | Inner_offset |
|---|---|---|---|---|---|---|---|---|---|---|
| chr1 | 230769615 | 230769683 | D1S1656 | 4 | CCTA [TCTA]n TCA [TCTA]n | 68 | 17 | 0.25 | 2 | 0 |
| chr2 | 218014858 | 218014950 | D2S1338 | 4 | [GGAA]n GGAC [GGAA]n [GGCA]n | 92 | 23 | 0.25 | 2 | 0 |
| chr2 | 1489652 | 1489684 | TPOX | 4 | [AATG]n | 32 | 8 | 0.25 | 2 | 0 |
| chr2 | 68011946 | 68011994 | D2S441 | 4 | [TCTA]n TCA [TCTA]n | 48 | 12 | 0.25 | 2 | 0 |
| chr3 | 45540738 | 45540802 | D3S1358 | 4 | [TCTA]n [TCTG]n [TCTA]n | 64 | 16 | 0.25 | 2 | 0 |
| chr4 | 154587735 | 154587823 | FGA | 4 | [GGAA]n GGAG [AAAG]n AGAA AAAA [GAAA]n | 88 | 22 | 0.25 | 2 | 0 |
| chr5 | 123775555 | 123775599 | D5S818 | 4 | [ATCT]n | 44 | 11 | 0.25 | 2 | 0 |
| chr5 | 150076323 | 150076375 | CSF1PO | 4 | [ATCT]n | 52 | 13 | 0.25 | 2 | 0 |
| chr6 | 88277143 | 88277245 | SE33 | 4 | [CTTT]n TT CT [CTTT]n | 102 | 25.2 | 0.25 | 2 | 0 |
| chr7 | 84160225 | 84160277 | D7S820 | 4 | [TATC]n | 52 | 13 | 0.25 | 2 | 0 |
| chr8 | 124894864 | 124894916 | D8S1179 | 4 | [TCTA]n [TCTG]n [TCTA]n | 52 | 13 | 0.25 | 2 | 0 |
| chr10 | 129294243 | 129294295 | D10S1248 | 4 | [GGAA]n | 52 | 13 | 0.25 | 2 | 0 |
| chr11 | 2171087 | 2171115 | TH01 | 4 | [AATG]n ATG [AATG]n | 28 | 7 | 0.25 | 2 | 0 |
| chr12 | 5983976 | 5984044 | vWA | 4 | [TAGA]n [CAGA]n TAGA | 68 | 17 | 0.25 | 2 | 0 |
| chr12 | 12297019 | 12297095 | D12S391 | 4 | [AGAT]n GA T [AGAT]n [AGAC]n AGAT | 76 | 19 | 0.25 | 2 | 0 |
| chr13 | 82148024 | 82148068 | D13S317 | 4 | [TATC]n | 44 | 11 | 0.25 | 2 | 0 |
| chr16 | 86352701 | 86352745 | D16S539 | 4 | [GATA]n | 44 | 11 | 0.25 | 2 | 0 |
| chr18 | 63281666 | 63281738 | D18S51 | 4 | [AGAA]n AG | 72 | 18 | 0.25 | 2 | 0 |
| chr19 | 29926234 | 29926298 | D19S433 | 4 | [CCTT]n ccta [CCTT]n cttt [CCTT]n | 64 | 14 | 0.25 | 2 | 8 |
| chr21 | 19181972 | 19182099 | D21S11 | 4 | [TCTA]n [TCTG]n [TCTA]n ta [TCTA]n tca [TCTA]n tccata [TCTA]n TA [TCTA]n | 127 | 29 | 0.25 | 2 | 11 |
| chr22 | 37140286 | 37140337 | D22S1045 | 3 | [ATT]n ACT [ATT]n | 51 | 17 | 0.25 | 2 | 0 |

The core loci were from the previous publication (Hares 2015).

**Table S2. Statistical summary of TRcaller called alleles from simulated read datasets**

| Coverage | Number of simulations | Number of alleles | Standard error | Accuracy | Missing ratio | Read type |
|---|---|---|---|---|---|---|
| 5 | 101 | 28.04 | 0.2256 | 0.701 | 0.299 | PE150 |
| 10 | 101 | 36.79 | 0.1531 | 0.9198 | 0.0802 | PE150 |
| 15 | 101 | 38.44 | 0.0959 | 0.9609 | 0.0391 | PE150 |
| 20 | 101 | 39.01 | 0.0838 | 0.9752 | 0.0248 | PE150 |
| 25 | 101 | 39.5 | 0.0591 | 0.9874 | 0.0126 | PE150 |
| 30 | 101 | 39.71 | 0.0514 | 0.9928 | 0.0072 | PE150 |
| 40 | 101 | 39.95 | 0.0217 | 0.9988 | 0.0012 | PE150 |
| 50 | 101 | 39.97 | 0.017 | 0.9993 | 0.0007 | PE150 |
| 60 | 101 | 40 | 0 | 1 | 0 | PE150 |
| 70 | 101 | 40 | 0 | 1 | 0 | PE150 |
| 80 | 101 | 40 | 0 | 1 | 0 | PE150 |
| 90 | 101 | 40 | 0 | 1 | 0 | PE150 |
| 100 | 101 | 40 | 0 | 1 | 0 | PE150 |
| 5 | 100 | 34.45 | 0.1956 | 0.8613 | 0.1388 | PE250 |
| 10 | 100 | 39.58 | 0.0572 | 0.9895 | 0.0105 | PE250 |
| 15 | 100 | 39.98 | 0.0141 | 0.9995 | 0.0005 | PE250 |
| 20 | 100 | 39.99 | 0.01 | 0.9998 | 0.0002 | PE250 |
| 25 | 100 | 40 | 0 | 1 | 0 | PE250 |
| 30 | 100 | 40 | 0 | 1 | 0 | PE250 |
| 40 | 100 | 40 | 0 | 1 | 0 | PE250 |
| 50 | 100 | 40 | 0 | 1 | 0 | PE250 |
| 60 | 100 | 40 | 0 | 1 | 0 | PE250 |
| 70 | 100 | 40 | 0 | 1 | 0 | PE250 |
| 80 | 100 | 40 | 0 | 1 | 0 | PE250 |
| 90 | 100 | 40 | 0 | 1 | 0 | PE250 |
| 100 | 100 | 40 | 0 | 1 | 0 | PE250 |
| 5 | 100 | 19.96 | 0.0197 | 0.998 | 0.002 | PLCCS |
| 10 | 100 | 20 | 0 | 1 | 0 | PLCCS |
| 15 | 100 | 20 | 0 | 1 | 0 | PLCCS |

| | | | | | | |
|---|---|---|---|---|---|---|
| 20 | 100 | 20 | 0 | 1 | 0 | PLCCS |
| 25 | 100 | 20 | 0 | 1 | 0 | PLCCS |
| 30 | 100 | 20 | 0 | 1 | 0 | PLCCS |
| 40 | 100 | 20 | 0 | 1 | 0 | PLCCS |
| 50 | 100 | 20 | 0 | 1 | 0 | PLCCS |
| 60 | 100 | 20 | 0 | 1 | 0 | PLCCS |
| 70 | 100 | 20 | 0 | 1 | 0 | PLCCS |
| 80 | 100 | 20 | 0 | 1 | 0 | PLCCS |
| 90 | 100 | 20 | 0 | 1 | 0 | PLCCS |
| 100 | 100 | 20 | 0 | 1 | 0 | PLCCS |

Note: The total expected number of alleles is 40. Accuracy was calculated by the number of called correct alleles divided by the expected number of alleles. No incorrect allele was found.

**Table S3**. ***Statistical summary of TRcaller called marker loci from simulated read datasets***

| Coverage | Number of simulations | Number of loci | Standard error | Accuracy | Missing ratio | Read type |
|---:|---:|---:|---:|---:|---:|---:|
| 5 | 101 | 17.554 | 0.113 | 0.878 | 0.122 | PE150 |
| 10 | 101 | 19.436 | 0.055 | 0.972 | 0.028 | PE150 |
| 15 | 101 | 19.683 | 0.047 | 0.984 | 0.016 | PE150 |
| 20 | 101 | 19.802 | 0.04 | 0.99 | 0.01 | PE150 |
| 25 | 101 | 19.95 | 0.022 | 0.998 | 0.002 | PE150 |
| 30 | 101 | 19.97 | 0.017 | 0.999 | 0.001 | PE150 |
| 40 | 101 | 20 | 0 | 1 | 0 | PE150 |
| 50 | 101 | 20 | 0 | 1 | 0 | PE150 |
| 60 | 101 | 20 | 0 | 1 | 0 | PE150 |
| 70 | 101 | 20 | 0 | 1 | 0 | PE150 |
| 80 | 101 | 20 | 0 | 1 | 0 | PE150 |
| 90 | 101 | 20 | 0 | 1 | 0 | PE150 |
| 100 | 101 | 20 | 0 | 1 | 0 | PE150 |
| 5 | 100 | 19.48 | 0.069 | 0.974 | 0.026 | PE250 |
| 10 | 100 | 20 | 0 | 1 | 0 | PE250 |
| 15 | 100 | 20 | 0 | 1 | 0 | PE250 |
| 20 | 100 | 20 | 0 | 1 | 0 | PE250 |
| 25 | 100 | 20 | 0 | 1 | 0 | PE250 |
| 30 | 100 | 20 | 0 | 1 | 0 | PE250 |
| 40 | 100 | 20 | 0 | 1 | 0 | PE250 |
| 50 | 100 | 20 | 0 | 1 | 0 | PE250 |
| 60 | 100 | 20 | 0 | 1 | 0 | PE250 |
| 70 | 100 | 20 | 0 | 1 | 0 | PE250 |
| 80 | 100 | 20 | 0 | 1 | 0 | PE250 |
| 90 | 100 | 20 | 0 | 1 | 0 | PE250 |
| 100 | 100 | 20 | 0 | 1 | 0 | PE250 |
| 5 | 100 | 19.96 | 0.02 | 0.998 | 0.002 | PLCCS |
| 10 | 100 | 20 | 0 | 1 | 0 | PLCCS |
| 15 | 100 | 20 | 0 | 1 | 0 | PLCCS |

| | | | | | | |
|---|---|---|---|---|---|---|
| 20 | 100 | 20 | 0 | 1 | 0 | PLCCS |
| 25 | 100 | 20 | 0 | 1 | 0 | PLCCS |
| 30 | 100 | 20 | 0 | 1 | 0 | PLCCS |
| 40 | 100 | 20 | 0 | 1 | 0 | PLCCS |
| 50 | 100 | 20 | 0 | 1 | 0 | PLCCS |
| 60 | 100 | 20 | 0 | 1 | 0 | PLCCS |
| 70 | 100 | 20 | 0 | 1 | 0 | PLCCS |
| 80 | 100 | 20 | 0 | 1 | 0 | PLCCS |
| 90 | 100 | 20 | 0 | 1 | 0 | PLCCS |
| 100 | 100 | 20 | 0 | 1 | 0 | PLCCS |

Note: The total expected number of marker loci are 20. Accuracy was calculated by the number of called loci divided by the expected number of loci.

**Table S4. The comparison of STR alleles between published data from Aalbers et al. 2020 and HipSTR**

| | |
|---|---|
| File name | Weir_data.hipSTR.comp.Statistic.xlsx |
| This file only lists the inconsistent alleles only | |

**Table S5. The comparison of STR alleles between published data from Aalbers et al. 2020 and TRcaller**

| | |
|---|---|
| File name | 289_samples_error_STRcaller_Weirs_manual_checked.xlsx |
| This file only lists the inconsistent alleles only | |

### Table S6. Links to publicly available sequencing datasets used in this study

| No. | Dataset | Link |
|---|---|---|
| 1 | Illumina NovoSeq WGS 2x150bp 300X per individual | ftp://ftp-trace.ncbi.nlm.nih.gov/ReferenceSamples/giab/data/AshkenazimTrio/HG002_NA24385_son/ NIST_HiSeq_HG002_Homogeneity-10953946/NHGRI_Illumina300X_AJtrio_novoalign_bams/HG002.GRCh38.300x.bam |
| 2 | Illumina NovoSeq WGS 2X250bp | ftp://ftp-trace.ncbi.nlm.nih.gov/ReferenceSamples/giab/data/AshkenazimTrio/HG002_NA24385_son/NIST_Illumina_2x250bps/ novoalign_bams/HG002.GRCh38.2x250.bam |
| 3 | 10X Genomics | ftp://ftp-trace.ncbi.nlm.nih.gov/ReferenceSamples/giab/data/AshkenazimTrio/HG002_NA24385_son/ 10XGenomics/NA24385_phased_possorted_bam.bam |
| 4 | PacBio Sequel II CCS 15kb_20kb chemistry2 | ftp://ftp-trace.ncbi.nlm.nih.gov/ReferenceSamples/giab/data/AshkenazimTrio/HG002_NA24385_son/PacBio_CCS_15kb_20kb_chemistry2/GRCh38/ HG002.SequelII.merged_15kb_20kb.pbmm2.GRCh38.haplotag.10x.bam |
| 5 | Oxford Nanopore MinION R10.4 | aws s3 sync --no-sign-request s3://ont-open-data/gm24385_q20_2021.10/ #data run: 5C, 5D and 5G |
| 6 | Disease PacBio, No-amp | https://downloads.pacbcloud.com/public/dataset/RepeatExpansionDisorders_NoAmp/analysis/align , samples bc1015-1021 |
| 7 | 1000 genome project | https://www.internationalgenome.org/data-portal/data-collection/30x-grch38 (reference ftp://ftp.1000genomes.ebi.ac.uk/vol1/ftp/technical/reference/GRCh38_reference_genome/GRCh38_full_analysis_set_plus_decoy_hla.fa ) |

**Table S7. Configuration information of 4 disease-associated TR loci**

| Chrom | ChromStart | ChromEnd | Name | Basic_motif_period | Motif | Ref_hap_length | Ref_allele | Stutter_ratio_threshold | Ploidy | Inner_offset |
|---|---|---|---|---|---|---|---|---|---|---|
| chr4 | 3076604 | 3076694 | HTT | 3 | [CAG]n | 90 | 30 | 0 | 2 | 0 |
| chr9 | 27573521 | 27573545 | C9orf72 | 6 | [GGGGCC]n | 24 | 4 | 0 | 2 | 0 |
| chrX | 146993569 | 146993629 | FMR1 | 3 | [CGG]n | 60 | 20 | 0 | 2 | 0 |
| chr22 | 46191235 | 46191305 | ATXN10 | 5 | [ATTGT]n | 70 | 14 | 0 | 2 | 0 |

**Table S8. The previously identified alleles from disease-associated tandem repeat loci**

| | |
|---|---|
| Samples | bc1015-bc1021 |
| Link | https://downloads.pacbcloud.com/public/dataset/RepeatExpansionDisorders_NoAmp/analysis/reports/ |
| Tools | https://github.com/PacificBiosciences/apps-scripts/tree/master/RepeatAnalysisTools |
